# Genetic risk for schizophrenia and developmental delay is associated with shape and microstructure of midline white matter structures

**DOI:** 10.1101/318238

**Authors:** Mark Drakesmith, Greg D Parker, Jacqueline Smith, Stefanie C Linden, Elliott Rees, Nigel Williams, Micheal J Owen, Marianne Van Den Bree, Jeremy Hall, Derek K Jones, David E J Linden

## Abstract

Genomic copy number variants (CNVs) are amongst the most highly penetrant genetic risk factors for neuropsychiatric disorders. The scarcity of carriers of individual CNVs and their phenotypical heterogeneity limits investigations of the associated neural mechanisms and endophenotypes. We applied a novel design based on CNV penetrance for schizophrenia and developmental delay that allows us to identify structural sequelae that are most relevant to neuropsychiatric disorders. Our focus on brain structural abnormalities was based on the hypothesis that convergent mechanisms contributing to neurodevelopmental disorders would likely manifest in the macro- and microstructure of white matter and cortical and subcortical grey matter. 21 adult participants carrying neuropsychiatric risk CNVs (including those located at 22q11.2, 15q11.2, 1q21.1, 16p11.2, and 17q12) and 15 age- and gender matched controls underwent T1-weighted structural, diffusion and quantitative T1 relaxometry MRI.

The macro- and microstructural properties of the cingulum bundles were associated with penetrance for both developmental delay and schizophrenia, in particular curvature along the anterior-posterior axis (Sz: p_corr_=0.026; DD: p_corr_=0.035) and intracellular volume fraction (Sz: p_corr_=0.019; DD: p_corr_=0.064) Further principal component analysis showed alterations in the interrelationships between the volumes of several mid-line white matter structures (Sz: p_corr_=0.055; DD, p_corr_=0.027). In particular, the ratio of volumes in the splenium and body of the corpus callosum was significantly associated with both penetrance scores (Sz: p=0.037; DD; p=0.006). Our results are consistent with the notion that a significant alteration in developmental trajectories of mid-line white-matter structures constitutes a common neurodevelopmental aberration contributing to risk for schizophrenia and intellectual disability.

## Introduction

Genetic variants are associated with neurodevelopmental disorders across conventional diagnostic classifications. This is evidenced both by common variants with low penetrance and by rarer but highly penetrant copy number variants (CNVs) ^1,2^. The mechanisms through which these genetic variants affect brain and behaviour are poorly understood, but genetic imaging studies in people selected for polygenic risk ^3^ or CNV carriers ^4^ can elucidate changes in brain development, structure and function that are not confounded by secondary disease effects. Because of their high penetrance, CNVs are particularly suited to translational studies elucidating disease mechanisms. However, rarity of CNVs makes it difficult to investigate their biological effects in humans. For most of the pathogenic CNVs identified for schizophrenia and developmental delay there is no information about neuroimaging correlates beyond small case series. Most neuroimaging studies of these CNVs have assessed effects of the 22q11.2 deletion (reviewed in ^5–7^). Neuroimaging studies in carriers of other CNVs have revealed highly heterogeneous changes in brain structure across CNVs (Table 1). Most finding relate to brain morphology (measured using regional volumes, cortical thickness, surface area etc.), with few studies reporting microstructural difference, usually measured with diffusion tensor metrics such as fractional anisotropy (FA) and mean diffusivity (MD). Although some of these findings corroborate those for non-CNV neuropsychiatric patients (e.g. structural correlates in 15q11.2 deletion carriers are similar to those found in dyslexia ^8^), in most cases it is difficult to corroborate findings in CNV patients with particular clinical features.

**Table 1.** Summary of neuroimaging findings in targeted CNVs. Literature found using search terms “<CNV> imaging” and “<CNV> MRI”. Findings based on single case studies or non-quantitative case series are italicised. Abbreviations:

| CNV (hg19) | Neuroimaging finding |  |
| --- | --- | --- |
|  | Morphological | Microstructural |
| 15q11.2 BP1-2 deletion (chr15:22,805,313-23,094,530) | Decreased volume in fusiform gyrus<sup>8</sup> | None |
| 15q13.3 BP4-5 duplication (chr15:31,080,645-32,462,776) | Abnormal bilateral hippocampal structure<sup>9</sup> | None |
| 15q13.3 BP4-5 deletion (chr15:31,080,645-32,462,776) | *Pachygyria and subcortical band heterotopia*<sup>10</sup> | -None |
| 16p11.2 distal duplication (chr16:28,823,196-29,046,783) | *cortical atrophy, thinning of corpus callosum, ventricular enlargement.*<sup>11</sup> |  |
| 16p11.2 deletion chr16:29,650,840-30,200,773 | Increased brain size. Increased cortical grey matter<sup>12</sup> Medio-dorsal thalamus, insula, ventral striatum, orbito-frontal cortex and fronto-striatal white matter<sup>13</sup> | Changes in fractional anisotropy and mean diffusivity in reward and language pathways; striatum, middle and superior temporal gyrus<sup>13</sup>. increased axial diffusivity in many major white matter tracts, including the anterior corpus callosum, internal and external capsules. Decreases in fibre orientation dispersion<sup>14</sup> |
| 17q12 duplication (chr17:34,815,904-36,217,432) | *Brainstem atrophy and hippocampal asymmetry*<sup>15</sup> | None |
| 1q21.1 deletion (chr1:146,527,987-147,394,444) | *No changes found*<sup>16</sup> | None |
| 1q21.1 duplication (chr1:146,527,987-147,394,444) | *Reduced corpus callosum volume, enlarged ventricles.*<sup>16</sup> | None |
| 22q11.2 deletion (chr22:19,037,332-21,466,726) | Whole-brain volumetric reductions, particularly in midline regions<sup>17-19</sup>. Increased cortical thickness<sup>20-22</sup>. Rostrocaudal gradient of volume reduction<sup>23</sup>. More comprehensive reviews are found in<sup>5-7</sup>. | Differences in fractional anisotropy and diffusivity parameters, e.g. lower fractional anisotropy in cingulum bundle and lower mean diffusivity in inferior longitudinal fasciculus<sup>24</sup> increase in fractional anisotropy in left inferior frontooccipital fasciculus, and a decrease in RD in right IFOF; increase in right hemisphere FA and a decrease in right RD in right cingulum; increase FA decrease RD within the right thalamo-frontal tract; decrease radial diffusivity in right inferior longitudinal fasciculus.<sup>21</sup> |
| 22q11.2 duplication (chr22:19,037,332-21,466,726) | Greater overall grey and white matter volumes and cortical surface area. Reduced cortical thickness. larger right hippocampus smaller caudate and corpus callosum volume (opposite findings compared to 22q11.2del)<sup>21</sup> | None |
| 3q29 deletion (chr3:195,720,167-197,354,826) | None | None |
| NRXN1 deletion (chr2:50145643-51259674) | *General lack of findings*<sup>25</sup> except for one case of hippocampal atrophy<sup>26</sup> | None |

Table 1 attests to the difficulty of collecting substantial quantitative data and drawing consistent conclusions about morphological and microstructural brain alterations in carriers from the literature, particularly of the rarer CNVs. We therefore conducted an analysis across a cohort of carriers of different CNVs, looking for features that correlate with the propensity of these CNVs to contribute to neuropsychiatric illness. We adopt a novel approach to characterising brain features in high-risk CNV carriers using penetrance scores previously calculated from a large cohort of neuropsychiatric CNV patients ^27^. This approach allows us to take account of the degree of pathogenicity of a genetic variant, testing the hypothesis that clinically more penetrant variants will also be associated with more salient neurodevelopmental changes as detected on neuroimaging. This method has the advantage over single-CNV studies that it enables the detection of convergent pathways common to several genetic variants, which are putatively most directly related to the pathophysiology of the associated diseases.

We explore structural brain phenotypes derived from neuroimaging data. These relate to macroscopic structure of cortex, including volume, surface area and cortical thickness, and the size of subcortical regions using T1-weighted structural MRI. We also quantify indices of white matter microstructure and morphology using diffusion MRI. We quantify tract volume and tract shape using a novel approach that extracts principal modes of feature variation of streamline shape^28^. We also quantify various microstructural parameters within the principal white matter pathways using metrics derived from diffusion tensor imaging (DTI), measures of axon density and dispersion using the neurite orientation dispersion and density imaging (NODDI)^29^ method and T1 relaxometry which provides a putative index of myelination ^30^.

In addition to examining each of these features separately, we also use principal component analysis (PCA) to identify any components across all these neuroanatomical features that show a strong effect of disease penetrance. The advantage of this approach is that it can highlight any prevalent components across correlated features that are not necessarily apparent when examining individual features separately.

## Methods

### Participants

The study was approved by the South Wales Research Ethics Committee and the Cardiff University School of Psychology Ethics Committee. All participants provided written informed consent.

MRI data were obtained from 21 CNV carriers and 15 controls. CNVs (Table 2) were targeted for their association with the development of schizophrenia and developmental disorder. Patients were recruited from NHS genetics clinics within the UK and through information disseminated by relevant support groups to their members. Full demographic details and clinical features for each patient are provided in the Supplementary Information (SI). Exclusion criteria included any contraindication to MRI.

**Table 2.**
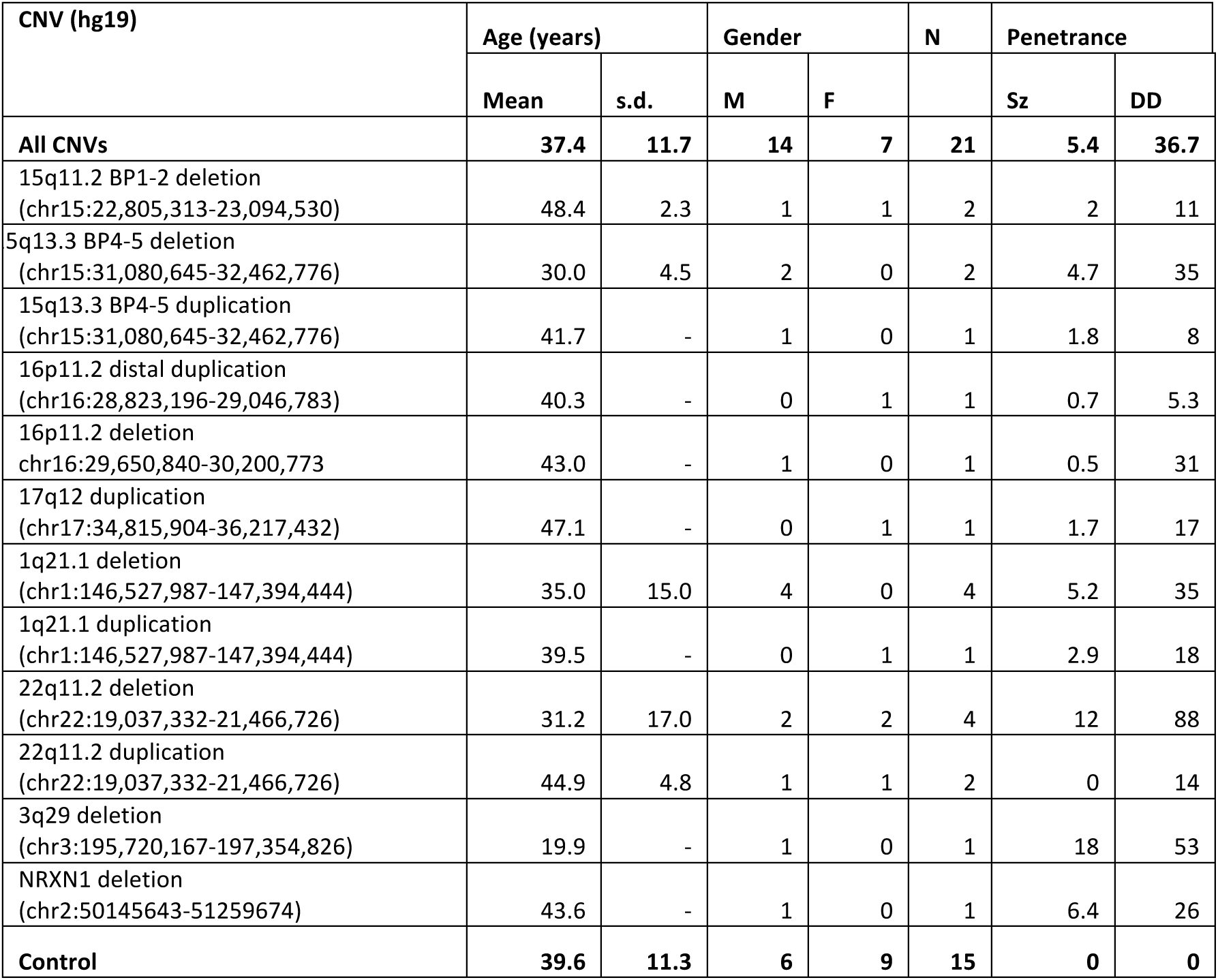
Demographic data for CNV and control

| CNV (hg19) | Age (years) |  | Gender |  | N | Penetrance |  |
| --- | --- | --- | --- | --- | --- | --- | --- |
|  | Mean | s.d. | M | F |  | Sz | DD |
| **All CNVs** | **37.4** | **11.7** | **14** | **7** | **21** | **5.4** | **36.7** |
| 15q11.2 BP1-2 deletion<br>(chr15:22,805,313-23,094,530) | 48.4 | 2.3 | 1 | 1 | 2 | 2 | 11 |
| 5q13.3 BP4-5 deletion<br>(chr15:31,080,645-32,462,776) | 30.0 | 4.5 | 2 | 0 | 2 | 4.7 | 35 |
| 15q13.3 BP4-5 duplication<br>(chr15:31,080,645-32,462,776) | 41.7 | - | 1 | 0 | 1 | 1.8 | 8 |
| 16p11.2 distal duplication<br>(chr16:28,823,196-29,046,783) | 40.3 | - | 0 | 1 | 1 | 0.7 | 5.3 |
| 16p11.2 deletion<br>chr16:29,650,840-30,200,773 | 43.0 | - | 1 | 0 | 1 | 0.5 | 31 |
| 17q12 duplication<br>(chr17:34,815,904-36,217,432) | 47.1 | - | 0 | 1 | 1 | 1.7 | 17 |
| 1q21.1 deletion<br>(chr1:146,527,987-147,394,444) | 35.0 | 15.0 | 4 | 0 | 4 | 5.2 | 35 |
| 1q21.1 duplication<br>(chr1:146,527,987-147,394,444) | 39.5 | - | 0 | 1 | 1 | 2.9 | 18 |
| 22q11.2 deletion<br>(chr22:19,037,332-21,466,726) | 31.2 | 17.0 | 2 | 2 | 4 | 12 | 88 |
| 22q11.2 duplication<br>(chr22:19,037,332-21,466,726) | 44.9 | 4.8 | 1 | 1 | 2 | 0 | 14 |
| 3q29 deletion<br>(chr3:195,720,167-197,354,826) | 19.9 | - | 1 | 0 | 1 | 18 | 53 |
| NRXN1 deletion<br>(chr2:50145643-51259674) | 43.6 | - | 1 | 0 | 1 | 6.4 | 26 |
| **Control** | **39.6** | **11.3** | **6** | **9** | **15** | **0** | **0** |

Controls were recruited via a local panel of volunteers. Control participants were chosen to match the age and gender of the CNV patients where possible. Criteria for inclusion were having no history of neurological or psychiatric disorders in addition to the screening for patients.

### Genotyping

To confirm CNV status, all patients and controls were genotyped using the Illumina HumanCoreExome whole genome SNP array that contains an additional 27,000 genetic variants at loci that had been previously implicated in neurological and psychiatric disease, which included CNVs. After processing the raw intensity data using Illumina Genome Studio software (version 2011.1), log R ratios and B allele frequencies were used to call CNVs using PennCNV (version 1.0.3) ^31^. CNVs were called if they spanned at least 10 informative SNPs and were joined if the distance between them was less than 50% of their combined length. CNVs were excluded if they were less than 10 kb in size, overlapped low copy repeats by more than 50% of their length, or had a probe density of less than 1 probe/20 kb. The Log R ratio and B allele frequency plots at each of the genomic regions of interest (chr1:146,527,987–147,394,444, chr2:50145643–51259674, chr3:195,720,167–197,354,826, chr15:22,805,313–23,094,530, chr15:31,080,645–32,462,776, chr16:28,823,196–29,046,783, chr16:29,650,840–30,200,773, chr17:29,107,491–30,265,075, chr22:19,037,332–21,466,726) were also manually inspected in order to confirm the presence of the CNV.

### MRI acquisition

All MRI data were acquired on a 3T General Electric HDx MRI system (GE Medical Systems, Milwaukee, WI) using an eight-channel receive-only head RF coil.

*<u>Structural</u>:* T1-weighted structural images were acquired with a 3D fast spoiled gradient echo (FSPGR) sequence (TR=7.8ms, TE=3.0ms, voxel size=1mm^3^ isomorphic).

*<u>Diffusion</u>*: A cardiac-gated diffusion-weighted spin-echo echo-planar imaging sequence was used to acquire high angular resolution diffusion weighted images (HARDI) ^32^. 60 gradient orientations at b=2000 s/mm^2^, and 30 directions at b=1200 s/mm^2,^ and 6 unweighted (b=0 s/mm^2^) images were acquired with the following parameters: TE=87 ms, 60 slices, slice thickness=2.4mm, FoV = 230×230 mm, Acquisition matrix = 96×96, resulting in data acquired with a a 2.4×2.4×2.4mm isotropic resolution. This was followed by zero-filling to a 128×128, in-plane matrix for the fast Fourier transform. The final image resolution was therefore 1.8×1.8×2.4mm.

*<u>Relaxometry</u>:* Relaxometry images were acquired using driven equilibrium single pulse observation of T1 with high-speed incorporation of RF field inhomogeneities (DESPOT1-HIFI) ^30^. A series of spoiled gradient echo (SPGR) images was acquired with 8 flip angles plus an additional inversion-recovery (IR) SPGR image. All images had TE=2.11ms and TR=4.7ms. SPGR images were acquired with flip angles of 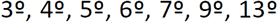 and 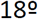. For the IR-SPGR acquisition, Inversion time = 450ms and flip angle = 5°.

### Grey-matter morphometry

Cortical reconstruction and volumetric segmentation were obtained from the T1-weighted structural images using Freesurfer (http://surfer.nmr.mgh.harvard.edu/) ^33^. The technical details of these procedures are described in prior publications ^34–40^. Grey matter was registered and parcelated to the Desikan-Killiany Atlas ^41^ (40 regions) and measures of volume, surface area, thickness and curvature were generated.

### Relaxometry pre-processing and analysis

Relaxometry data were pre-processed using FSL v5.0 ^42^. All SPGR/IR-SPGR images were coregistered to each other using a rigid affine transform and skull-stripped ^43^. Relaxation rate (R1 = 1/T1) maps were derived using Driven Equilibrium Single Pulse Observation of T1 with High-Speed Incorporation of RF Field Inhomogeneities (DESPOT1-HIFI)^30^, which incorporates correction for B1 field inhomogeneities with in-house code. A synthetic T1-weigted image was computed from the quantitative T1 map with contrast matching that of the FSPGR image. This was used as a reference for transforming the R1 (maps, which were then warped to the T1-weighted space.

### Diffusion MRI pre-processing

HARDI data were pre-processed in ExploreDTI v4.8.3 ^44^. Data were corrected for motion, eddy currents and field inhomogeneities prior to tractography. Motion artefacts and eddy current distortions were corrected with B-matrix rotation using the approach of ^45^. A comparison of subject motion between the two groups is shown in the supplementary materials.

Field inhomogeneities were corrected using the approach of ^46^. DWIs were non-linearly warped to the T1-weighted image using the fractional anisotropy map from the DWIs as a reference. Warps were computed using Elastix ^47^ using normalised mutual information cost function and constraining deformations to the phase-encoding direction. The corrected DWIs are therefore in the same (undistorted) space as the T1-weighted structural images.

### DTI analysis

The corrected HARDI data from the b=1200 s/mm^2^ shell were fitted to the diffusion tensor (DT) and corrected for CSF-partial volume effects ^48^ was applied to the DTs. The b=1200 s/mm^2^ shell was used as this is the domain in which the DT representation applies. The fractional anisotropy (FA), mean (MD), axial (AD) and radial (RD) diffusivities then computed from the DT. Intra-scan head motion was quantified and assessed for potential impact on subsequent statistics (see supplementary material, section 4).

### NODDI (neurite orientation dispersion and density imaging) analysis

NODDI ^29^ was performed using both the b=1200 s/mm^2^ and b=2000 s/mm^2^ diffusion shells using the NODDI toolbox v0.9 (http://mig.cs.ucl.ac.uk/mig/mig/index.php/?n=Tutorial.NODDImatlab/). NODDI yields three parameters that describe the microstructure of the tissue in each voxel, intracellular volume fraction (ICVF), isotropic fraction (ISOF), and orientation dispersion index (ODI).

### Tractography

Whole-brain tractography was performed using the damped Richardson-Lucy algorithm ^49^. This is a modified spherical deconvolution method which is more robust to spurious peaks in the fibre orientation distribution (FOD) than standard spherical deconvolution methods ^50^. RESDORE ^51^ was also applied to remove corrupted voxels from the FOD calculation. The tractography algorithm used is that of Basser et al., [2000] which uses a uniform step size. Seed points were arranged in a 3×3×3mm grid in white matter, step size = 1mm, angle threshold = 45°, length threshold = 20–500mm, FOD threshold = 0.05, *ϐ* = 1.77, *λ* = 0.0019, *η*=0.04, number of iterations = 200 (See ^49^ for full details of these parameters).

Additional anatomical constraints were introduced to ensure minimal contamination from spurious streamline trajectories through grey matter. A segmentation of the T1 weighted images was performed using FSL-FAST and was used to apply a mask to the streamlines, such that streamlines were forced to terminate when they entered grey matter. There were no explicit masking of CSF, however, the termination criteria used for the dRL algorithm, which is based on the amplitude of the FOD peak, ensures that no streamlines enter isotropic regions such as CSF.

### Tract segmentation and shape analysis

Tract shape analyses was performed to quantify the shapes of white matter pathways ^28^. Subject whole brain tractography results were affinely (registered to a standard MNI template (preserving shape but eliminating variance in position/orientation). Streamlines were then re-parameterised to 30 knot-points (spline interpolation), translated to the origin and reduced to a feature vector through coordinate concatenation. PCA was then applied to determine principal modes of feature vector variation and, by extension, variation of streamline shape. Following decomposition of the feature vectors onto the first seven PCA eigenvectors (representing a set of streamline shape basis functions encapsulating ~95% of observed shape variation), the resultant weight vectors were clustered (k-means, k=800) and, for each subject, histograms of streamline cluster membership recorded.

Streamlines were segmented into the following 19 tract bundles specific pathways, based on the descriptors computed from previously generated statistical models: Bilateral arcuate, uncinate, inferior longitudinal, superior longitudinal, fronto-occipital fasciculi, cingulum (dorsal and parahippocampal parts), fornix (left and right branches) and corpus callosum (splenium, body and genu). This yielded a total of 194 shape descriptors.

In addition to shape, volume of each bundle was also computed by counting voxels traversed by streamlines in each bundle and normalising to voxel volume.

All 9 microstructural measurements (from DTI, relaxometry and NODDI) were registered to streamline points in each bundle and the median value taken for each bundle. This yields a total of 190 (19 bundles x 10 variables) microstructural variables.

### CNV Penetrance scores

To assess the contribution of genetic loading for each CNV for psychopathology, and to accommodate the small sample size of each individual CNV within the cohort, penetrance cores were used for statistical analysis. The CNV penetrance scores used in the current study correspond to the probability of manifesting a given phenotype in carriers of that CNV. Estimates of CNV penetrance for the development of schizophrenia and ID/DD/ASD were obtained from Kirov et al 2014^27^. These authors used CNV data from large cohorts of patients diagnosed with schizophrenia, ID/DD/ASD and non-psychiatric controls, to estimate the rate of specific neuropsychiatric CNVs among these disease and non-disease populations, and then estimated CNV penetrance as the probability of carrying a specific CNV given disease status (i.e. rate of the CNV in schizophrenia or ID/DD/ASD), divided by the probability of carrying the CNV in the general population (which includes both case and control populations). Full details on how CNV penetrance scores were calculated can be found in ^27^.

### Statistical analysis

To assess the contribution of genetic loading for each CNV for psychopathology, and to accommodate the small sample size of each individual CNV within the cohort, each participant was assigned a penetrance score for schizophrenia and developmental delay, as previously computed in Kirov et al 2014 ^27^

A general linear model was applied to identify relationships between the penetrance scores and the imaging variable of interest. Age and gender were included as covariates. Furthermore, for volumetric measures, total brain volume was treated as a covariate and intra-scan head motion was included as a covariate for all diffusion-derived measures.

Multiple comparisons were corrected with permutation testing (5000 iterations) correcting across all observations within the same imaging domain, a correction that will also control for any distributions that are non-Gaussian. To verify effects are due to penetrance and not simply for the presence of CNVs, equivalent analysis was performed using only a binary classification of CNV carriers vs controls.

To identify multivariate features in the data, a principal component analysis (PCA) of all imaging variables was performed. The components were subject to the same statistical analysis as the variables.

## Results

### Relation between penetrance scores

There was a significant correlation between the penetrance scores P_Sz_ and P_DD_ (ρ= 0.77, p=7.7 × 10^−9^).

### White matter morphology

The shape analysis revealed significant alteration of one component of the left cingulum bundle that was significantly correlated with both P_Sz_ (t=4.195, p=2.11×10^−4^, p_corr_ = 0.026) and P_DD_ (t=4.06, p=3.1×10^−5^, p_corr_=0.035) (Figure 1). This component reflects the curvature of the dorsal cingulum along the AP axis (Figure 2) with higher penetrance associated with greater curvature of the dorsal cingulum. The same effect was seen in the corresponding component of the right dorsal cingulum with P_DD_ (t=-5.60, p=1.81×10^−6^, p_corr_ =8.00×10^−4^). A strong effect was also observed for P_Sz_, but this did not survive permutation correction (t=−3.601, p=0.001, p_corr_ =0.11). In the binary comparison of CNV carriers vs controls, these effects were non-significant (left cingulum: t=1.41, p=0.17; right cingulum: t=−1.94, p=0.062). Note that the sign of the descriptor from the analysis is arbitrary, so although the effect was in the opposite direction in the right compared to the left, the modes of variation also are also reversed between left and right. Therefore both left and right cingulum bundles are showing the same geometric variation - greater curvature associated with higher penetrance.

**Figure 1.**
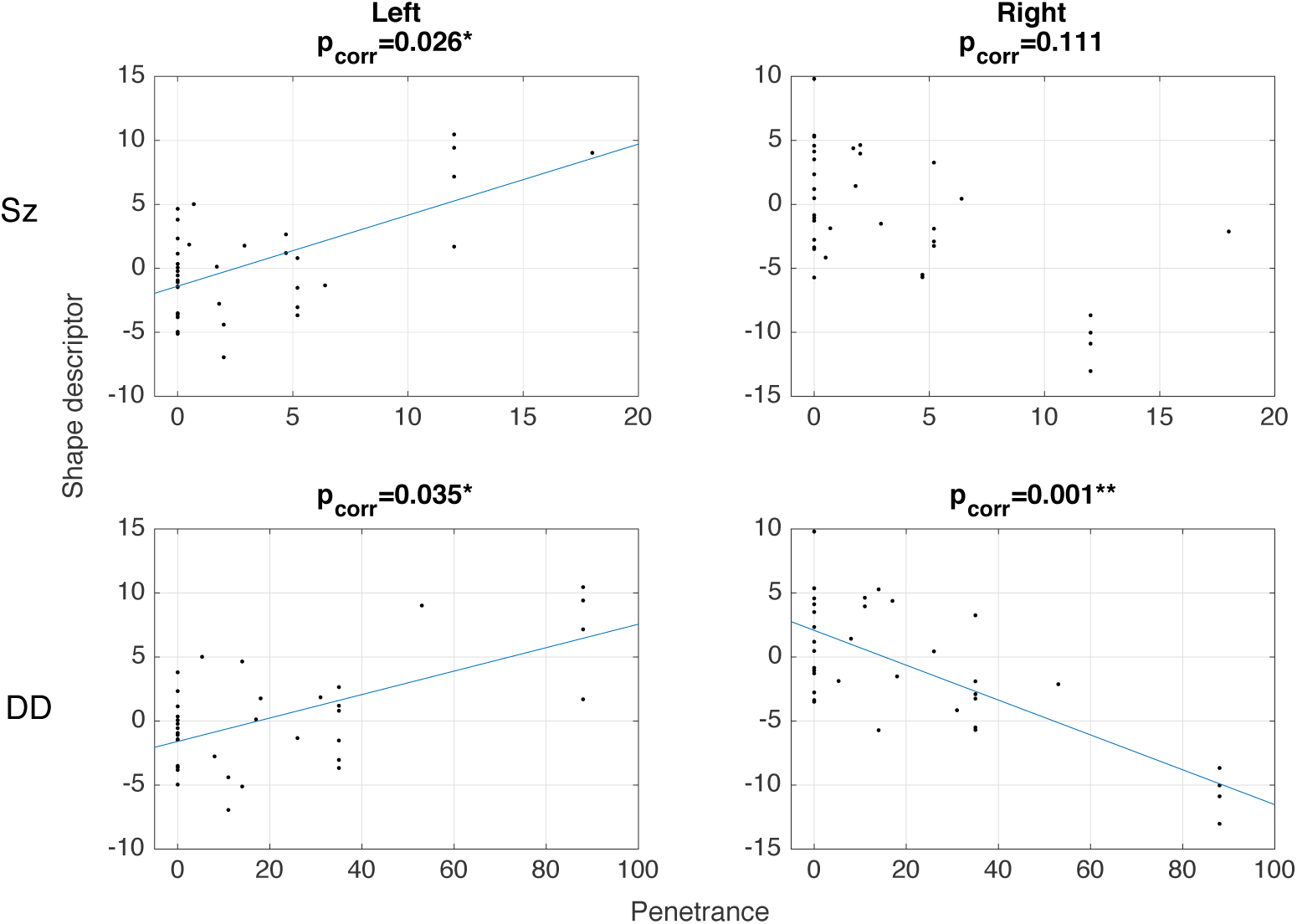
Scatterplots of the left and right homologous shape descriptors (visualised in **Figure 2**) in cingulum bundles against P_Sz_ and P_DD_, with associated regression lines (note the sign of the shape descriptor in the right cingulum was flipped for clarity).

**Figure 2.**
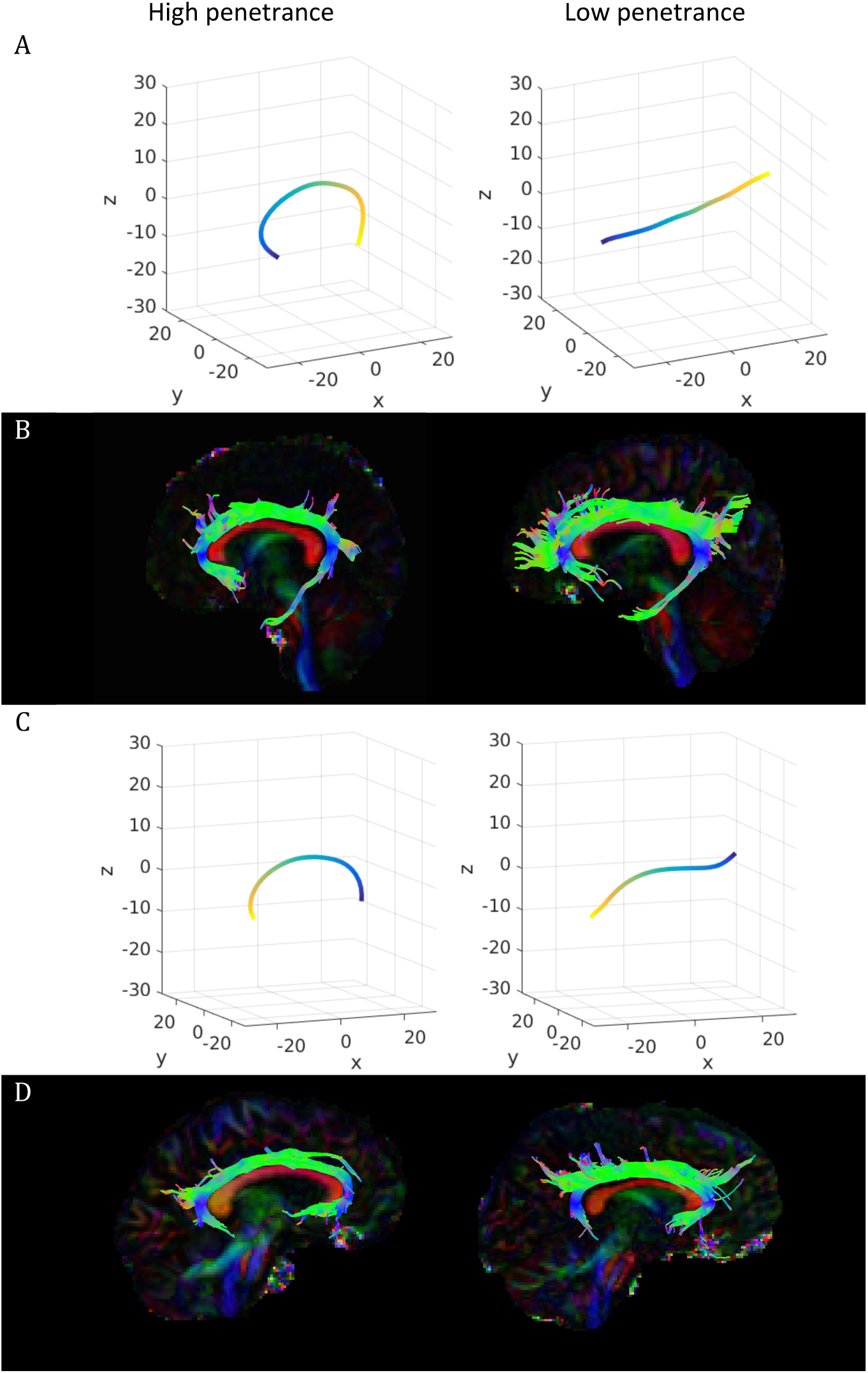
Shape descriptors and examples of segmented tracts for left cingulum (A-B) and right cingulum (C-D). Shape descriptors (A, C) show the mode of variation within the maximum (left) and minimum (right) range observed. Example tracts are shown for a patient with high P_DD_ (left) and a typical control (right).

**Figure 3.**
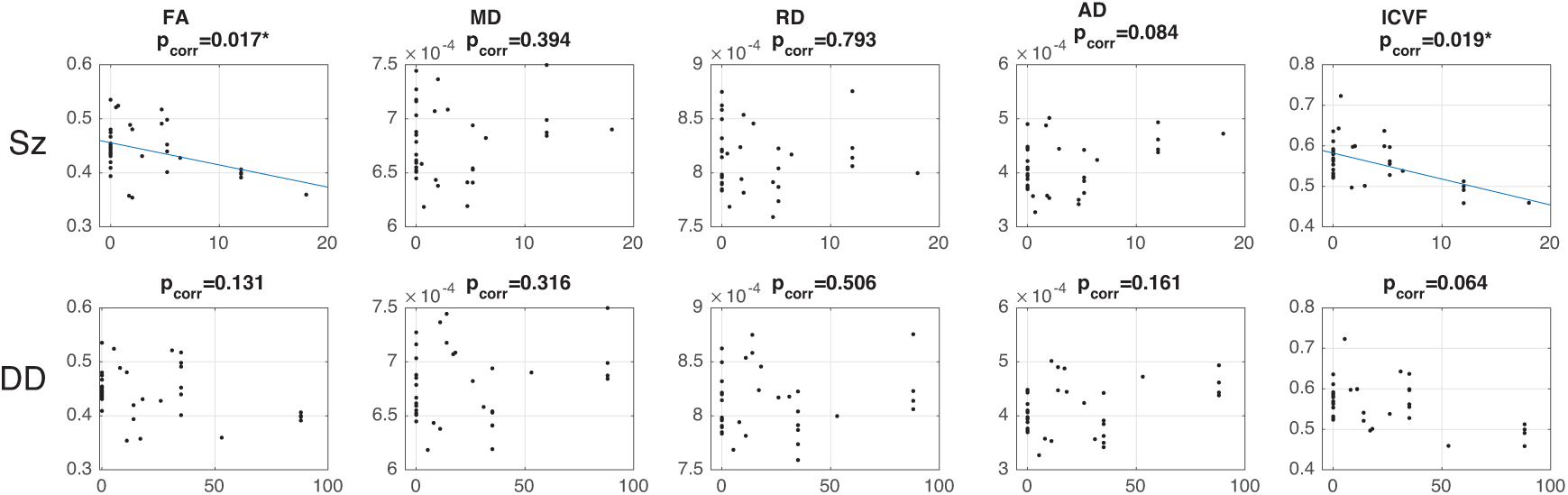
Scatterplots of various microstructural measures in the left cingulum against penetrance (Sz on top row, DD on bottom row). Linear regression fit lines are shown where effects are found to be significant or close-to-significant.

No other tract shape descriptors showed any significant effect although there were marginal effects of P_Sz_ on shape in the right parahippocampal cingulum and cortico-spinal tract. There were also marginal effects of P_DD_ on shape in the body of the corpus callosum. (see SI for full results).

Tract volumes show significant effects of P_Sz_ on the right corticospinal tract (t=−3.163, p=0.0035, p_corr_ =0.030) and a marginal effect on the right uncinate fasiculus (t=−2.881, p=0.0072, p_corr_ = 0.057). P_DD_ was associated with lower volumes of the right cingulum (t=−3.698, p=8.4×10^−4^, p_corr_=0.0094) and the right uncinate fasiculus (t=−3.617, p=0.0010, p_corr_=0.011). Right corticospinal tract (t=−2.605, p=0.014) and right uncinate fasiculus (t=−2.093, p=0.014) show some effects using the binary comparison, indicating significant group differences.

### White matter microstructure

There were significant correlations with both penetrance scores with ICVF in the left cingulum. There was also a significant association between P_Sz_ and FA. The findings are summarised in table 3. The general pattern is that penetrance scores are negatively correlated with FA and positively correlated with diffusivities, particularly axial diffusivity. No effects were observed for R1 (see SI for full results).

**Table 3.** Summary of statistics relating to DTI and NODDI measures of the left cingulum bundle. * significant at p<0.05 uncorrected, ** significant at p_corr_<0.05 (corrected).

| Method | Measure | $P_{Sz}$ | | | $P_{DD}$ | | | CNV carrier vs control | |
| --- | --- | --- | --- | --- | --- | --- | --- | --- | --- |
| | | t | p | $p_{corr}$ | t | p | $p_{corr}$ | t | p |
| DTI | FA | -3.290 | 0.002 | 0.017** | -2.316 | 0.027 | 0.131* | -1.291 | 0.352 |
|  | MD | 1.448 | 0.161 | 0.394 | 1.641 | 0.111 | 0.316 | 0.804 | 0.222 |
|  | RD | 0.620 | 0.545 | 0.793 | 1.232 | 0.227 | 0.506 | 0.897 | 0.243 |
|  | AD | 2.570 | 0.016 | 0.084* | 2.178 | 0.037 | 0.131* | 0.981 | 0.214 |
| NODDI | ICVF | -3.258 | 0.003 | 0.019** | -2.644 | 0.013 | 0.064* | -0.511 | 0.613 |
|  | ISOF | -2.551 | 0.016 | 0.072* | -1.509 | 0.141 | 0.370 | 0.535 | 0.596 |
|  | ODI | 1.905 | 0.066 | 0.342 | 1.403 | 0.171 | 0.577 | 1.010 | 0.320 |

Other microstructural metrics that were significant were a significant positive relationship between P_Sz_ and ODI in left and right inferior frontooccipital fasciculi (left: t=3.757, p=7.1×10^−4^, p_corr_=0.0072; right: t=4.034, p=3.3×10^−4^, p_corr_=0.0042).

### Other structural measures

There were no significant effects on grey matter morphometric or in gross brain morphometric measurements (see SI for full results).

### Principal component analysis

The first 34 components were tested for effects of penetrance. Only the 8^th^ largest component (PC8) was found to be significantly associated with P_DD_ (t=−3.033, p=0.005, p_corr_=0.027) and marginally significant for P_Sz_ (t=−2.560, p=0.016, p_corr_=0.055). This component shows no significant effect when tested with the binary CNV model (t=−2.431, p=0.021, p_corr_=0.324). This component is heavily weighted by variables relating to white matter volume (figure 4). The two largest anatomical regions with the highest weighting on this component are the body and the splenium of the corpus callosum. These areas were weighted in opposite directions, which would be compatible with the altered curvature observed for the risk scores (see above, Figure 1).

**Figure 4.**
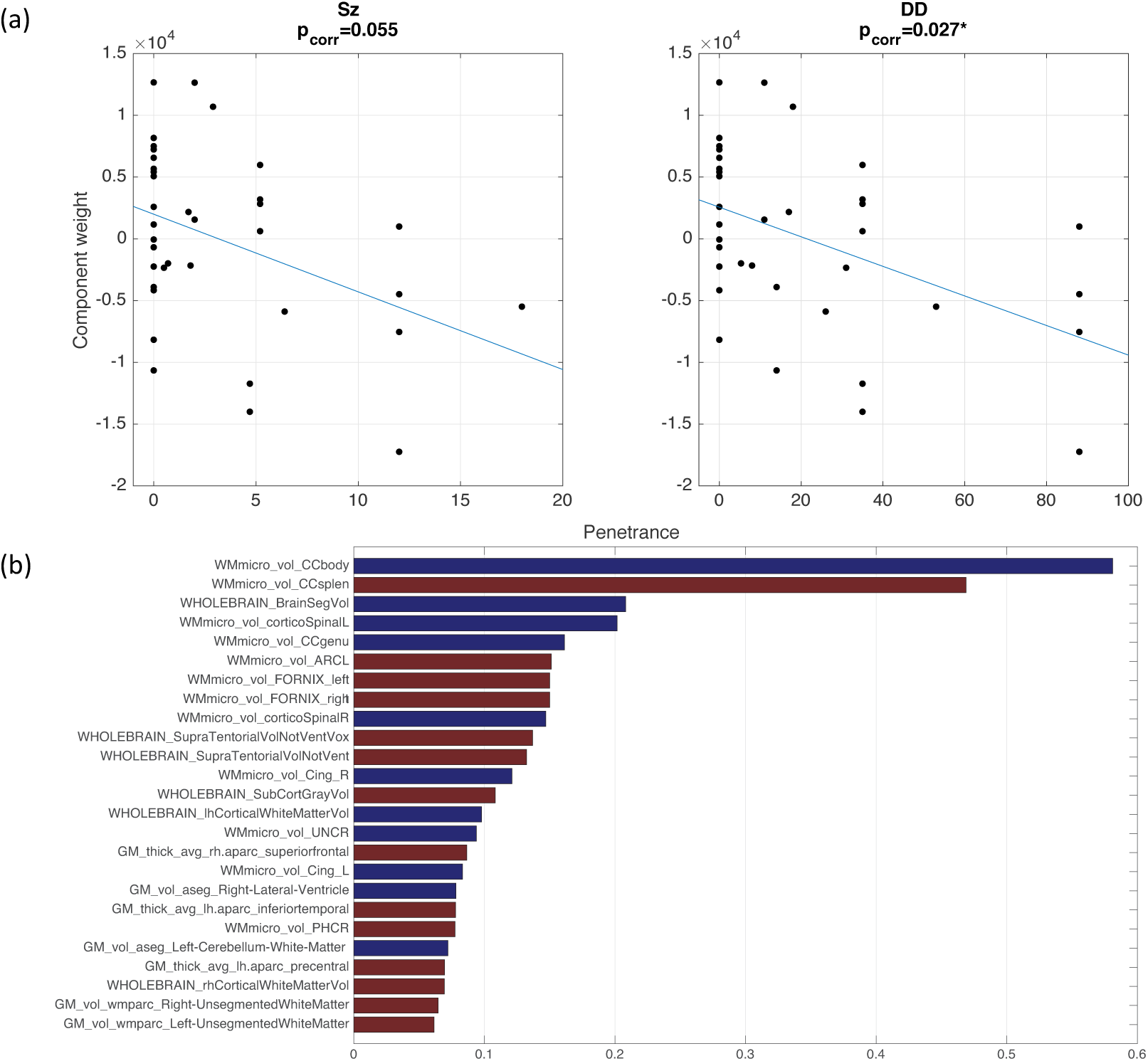
(a) Scatterplots of PC8 against P_Sz_ and P_DD_ (b) Weighting of each imaging variable in PC 8, showing the top 25 weightings. Blue bars indicate positive weighting, red bars indicate negative weighting.

Relative volumes of segments of the corpus callosum for extreme cases of PC8 weight are visualised in Figure 6. This shows the body is relatively smaller in high-penetrance cases (low PC8 weight) compared to low-penetrance cases (high PC8 weight). A post-hoc test was performed on the ratio of volumes between the CC body and splenium that confirms negative correlation with both penetrance scores (P_Sz_: t=−2.193, p=0.036; P_DD_; t=−2.931, p=0.006). The volumetric interrelationships between the white mater structural variables strongly represented in PC8 are visualised in Figure 5.

**Figure 5.**
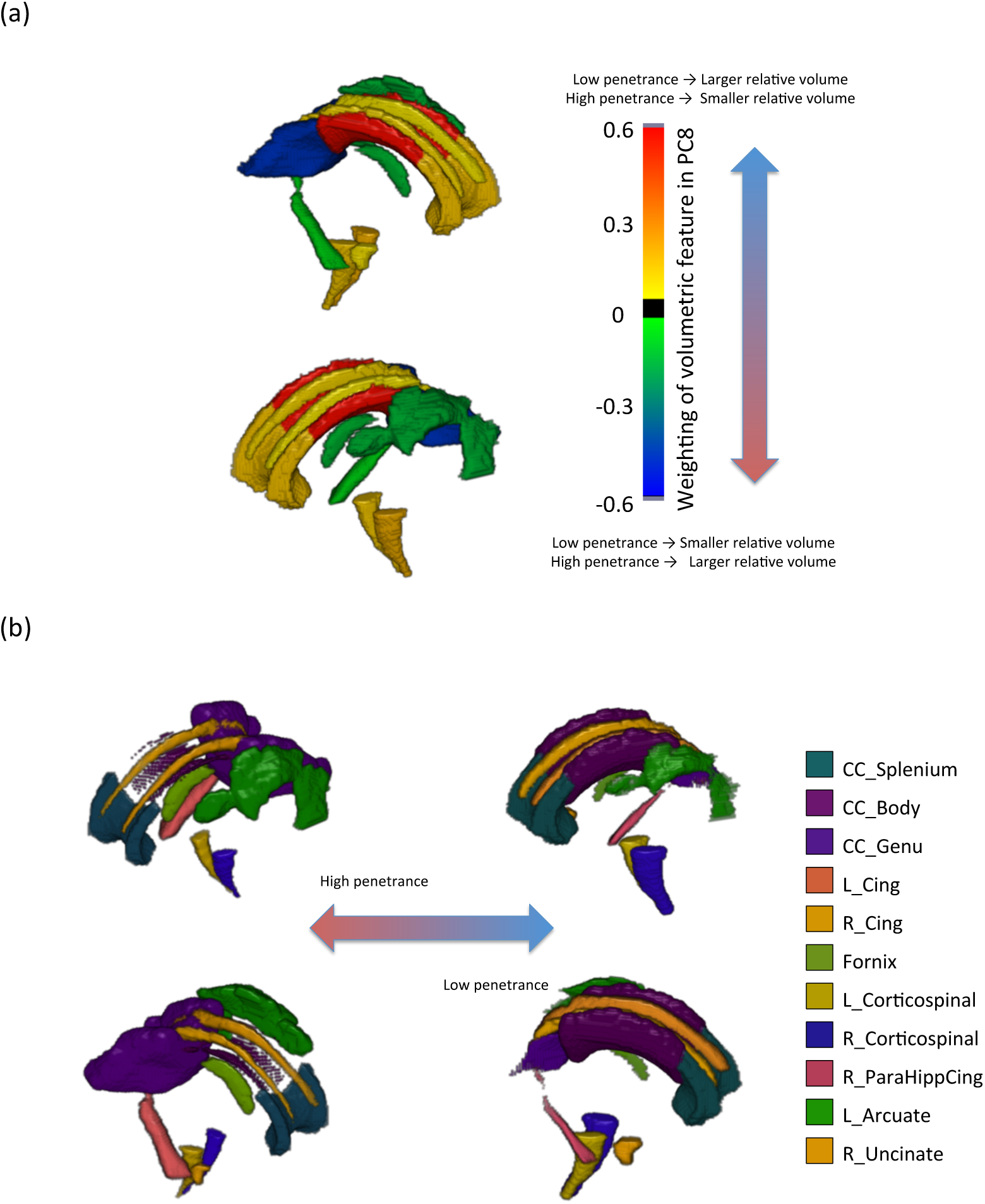
(a) Volumetric change associated with white matter structures strongly represented in PC8 rendered on the JHU atlas, with positive values corresponding to larger volumes for low penetrance (or smaller volume for higher penetrance) and negative values corresponding to smaller volumes for low penetrance (or larger volume for lower penetrance). (b) Volumetric interrelationships of relevant white matter volumes strongly associated with PC8. Volumes taken form the JHU atlas were inflated or eroded proportional to the weighting in PC8.

**Figure 6.**
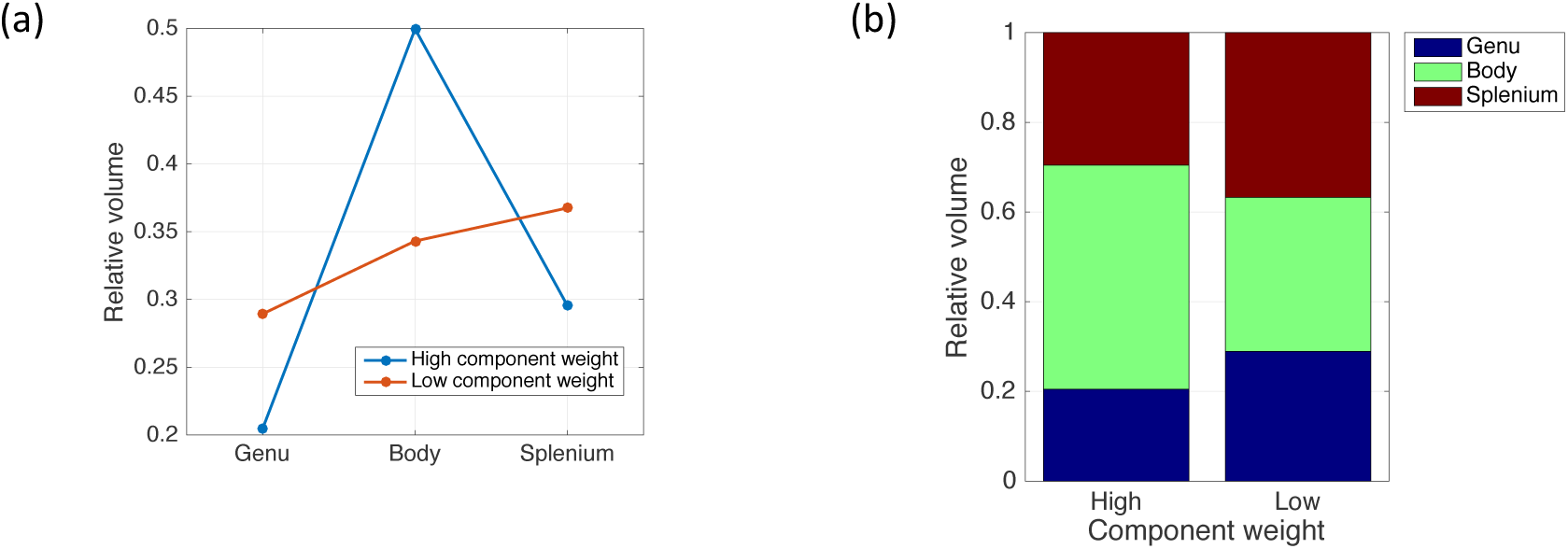
Relative volumes of 3 segments of corpus callosum for extreme cases of high and low component weight

## Discussion

In this study we investigated macrostructural / morphological and microstructural alterations associated with CNVs with high penetrance for schizophrenia and developmental delay. We utilised a novel approach to characterising morphology of white matter fibre bundles in combination with more traditional metrics of brain morphology and microstructure to identify features and principal components of these features that characterise neuropsychiatric CNVs. We found altered morphology in the left and right cingulum. We also found a significant reduction in FA and ICVF in this structure, and our preliminary interpretation of this result is that it reflects a reduction in axon density in this pathway rather than myelin because no effects of relaxometry measures were observed (although it should be noted this may be due to comparatively lower sensitivity of R1 compared to DTI metrics ^53^). There was also an overall positive trend of increased diffusivity with increased penetrance, which is also consistent with a decrease in fibre density in high penetrance individuals.

A converging finding comes from the PCA analysis, in which PC8 shows a strong effect of penetrance. This component is heavily weighted by midline white matter structures, in particular, the three components of the corpus callosum. Many of these volumes did not show effects when tested directly, which is likely due to discordant signs between the weighting of features within the component. Most prominently, the splenium has opposite weighting to the body and genu of the corpus callosum. This suggests that rather than there being absolute volumetric effects of penetrance, penetrance affects the volumetric interrelationships between these structures. The corpus callosum is of particular interest, because the altered inter-relationships in the three segments suggest alteration of the arrangement of fibres along the AP axis. This is consistent with the increase in curvature of the cingulum along the AP axis, which wraps dorsally around the corpus callosum. We obtained these findings in the absence of any effects of gross brain morphology. There is no apparent relationship between the overall brain size or shape and penetrance. Therefore, it appears that these high-risk CNVs lead to neurodevelopmental alterations that are associated with penetrance, but manifest more subtly than in gross brain morphology. Also of note is that the results of the binary model show weak effects for these metrics, adding further credence to the view that these features are related to CNV penetrance for neuropsychiatric illness rather than simply due to the presence of CNVs.

Altered brain development appears to manifest in the following way: there is increase in forces^54^ applied parallel to the AP axis. The distribution of axons along the length of the corpus callosum is altered, which causes a change in the forces applied to the cingulum during development. Some findings from 22q11.2 deletion patients appear to corroborate our findings, reporting a larger area of the mid-sagittal section of the corpus callosum, in particular the posterior part ^54–56^ which corroborates the pattern of fractional corpus callosum volumes observed in this study.

In terms of the mechanisms underlying the cingulum shape / microstructural abnormalities, the embryological literature indicates that the genes affected by the CNVs we have studied are expressed in the developing brain (e.g., for those in 22q11.2 see ^57^). It is therefore likely that alterations in brain morphology will start to occur at this very early stage of development. Abnormal brain morphology can impact on mechanical processes during embryonic brain development, leading to altered structure of the cingulum and consequently morphological differences in the arrangement of fibres along the corpus callosum, as suggested by mechanical models of morphogenesis ^58^. The early formation of neurons has been shown to guide the trajectory of axonal process ^59^. Therefore, we can speculate that early disturbances of head shape can lead to further disruption of axonal processes, causing a reduction in axon density. There is evidence for altered neuronal migration during brain development in both humans and mouse models of 22q11.2 deletion ^60^. Kates et al even speculate that 22q11.2 deletion is a disorder of axonal guidance as indicated by differential developmental trajectories between 22q11.2 deletion carriers and controls in microstructural measurements ^61,62^.

The changes in cingulum microstructure that we report are largely compatible with previous reports in schizophrenia ^63–65^. Similar finding have been observed in 1^st^ degree relatives of schizophrenia patients ^66^ and (for right cingulum) for carriers of the common schizophrenia risk variant at ZNF840A ^67^ and in a sample of adolescents and young adults with 22q11.2 deletion syndrome ^68^ where the most robust finding was AD reduction (without corresponding increases in RD) in the anterior and dorsal cingulum.

There is little discrimination between the effects of the two penetrance scores. All significant effects observed apply to both scores, with a few exceptions, for example, increased fibre dispersion, in bilateral inferior frontooccipital cortices associated with higher penetrance for schizophrenia. This is also evident in the high correlation between penetrance scores. This suggests that the neurodevelopmental indicators observed are not attributable to a particular psychopathology, but rather that these features reflect penetrance for neuropsychiatric illness more generally, a notion which has also been proposed for the clinical phenotype ^21^. However, it should be stressed, due to the comparatively weak effects seen in the binary model, this effect is not simply due to the presence of a CNV.

In summary, we reveal significant morphological and microstructural features associated with penetrance for neuropsychiatric illness in a cohort of CNV carriers. The most pronounced of these features is curvature of the cingulum bundle and volumetric interrelationships between difference segments of the corpus callosum. These features are consistent with a common neurodevelopmental trajectory, which does not manifest in gross brain morphological changes, but in more subtle alterations in the morphological interrelationships in mid-line white-matter structures. This can lead to downstream effects on cognitive and intellectual impairments that commonly arise in these CNVs.

## Acknowledgments

This study was funded by two Wellcome Trust Strategic Awards (100202/Z/12/Z, 104943/Z/14/Z) and a Wellcome Trust New Investigator awarded to DKJ (096646/Z/11/Z). We would also like to thank the members of the field team who assisted in recruitment, assessment and data collection, Ffion Evans, Kali Barawi, and Vera Schroeter.

